# Phenotypic Plasticity and Competition Shape Therapy Sequencing in HER2^+^/HER2^‒^ Breast Cancer: A Mathematical Framework

**DOI:** 10.64898/2025.12.27.696676

**Authors:** Aleksandra Gavrilova, Trachette Jackson, Nizhum Rahman

## Abstract

Tumour heterogeneity and phenotypic plasticity are major drivers of treatment failure in cancer, enabling rapid adaptation under therapeutic pressure. In HER2-positive (HER2^+^) breast cancer, tumours often contain both HER2^+^ and HER2-negative (HER2^−^) cells whose interactions complicate schedule design.

We develop a compact ordinary differential equation framework for intratumoral HER2^+^/^−^ dynamics that integrates phenotypic plasticity with density-dependent growth and inter-phenotype competition. Phenotype-specific therapies are incorporated through simple pharmacodynamic surrogates: Paclitaxel chemotherapy acting primarily on HER2^+^ cells and Notch-pathway inhibition targeting HER2^−^ cells. We use the model to compare staggered and simultaneous treatment schedules.

The results show that treatment order and relative intensity critically shape long-term tumour composition. Targeted-first schedules can exhibit competitive release, whereby subsequent aggressive chemotherapy unintentionally favours HER2^−^ expansion. In contrast, simultaneous initiation suppresses both phenotypes more effectively and avoids strong rebound. These findings highlight the importance of ecological structure in therapy design and support simultaneous combination therapy followed by targeted maintenance.

## 1 Introduction

Cancer remains a major public health burden. In the United States alone, an estimated 2,041,910 new cancer cases and 618,120 cancer deaths are expected in 2025 [1, 2]. Breast cancer is the most commonly diagnosed cancer, with an estimated 316,950 new invasive cases in women and 2,800 in men in 2025, and remains a leading cause of cancer mortality among women [1, 2]. Despite therapeutic advances, its population impact continues to grow, with notable increases in incidence documented over the last decade [3].

Molecular profiling has established that “breast cancer” comprises biologically distinct subtypes with different outcomes and treatment responses [4, 5]. Among these, HER2-positive (HER2+) disease is defined by amplification/overexpression of the ERBB2 (HER2) receptor and an aggressive clinical phenotype [5, 6]. HER2-targeted therapy (e.g., trastuzumab) transformed outcomes in both metastatic and early-stage settings [7, 8], and dual HER2 blockade (pertuzumab+trastuzumab+docetaxel) further improved overall survival [9, 10]. Nevertheless, resistance and relapse remain common, motivating a deeper mechanistic understanding and improved therapeutic strategies [11].

A central driver of therapy failure is intratumoral heterogeneity-the coexistence of genetically, phenotypically, and microenvironmentally diverse cell populations that enable rapid adaptation under treatment pressure [12–15]. Beyond fixed genetic diversity, *cell plasticity* allows tumor cells to transition between phenotypic states, replenishing therapy-tolerant compartments [16,17]. In HER2+ breast cancer, plasticity spans cancer stem-like, progenitor, and bulk-tumor states [18]. Elegant patient-derived models of circulating tumor cells (CTCs) demonstrated rapid, bidirectional interconversion between HER2+ and HER2-phenotypes within as few as four cell doublings, with therapy-tolerant states enriched under cytotoxic stress [19]. Phenotypic composition is further shaped by symmetric and asymmetric divisions, which regulate lineage balance and heterogeneity [20]. These ecological interactions can be frequency-dependent, making treatment outcomes sensitive to population structure and competitive dynamics [21].

The Notch pathway is tightly connected to these behaviors. Notch–HER2 bidirectional crosstalk modulates proliferation, stemness, and therapeutic tolerance in breast cancer [22, 23]. Notch activity is linked to dormancy and recurrence: it is upregulated after HER2 pathway inhibition, persists in residual/dormant cells, and its inhibition delays recurrence in preclinical models [24–26]. Collectively, these data suggest that coupling HER2-directed therapy with strategies that reshape phenotypic transitions and micro-ecological competition could extend durability of response.

**Our goal in this work** is to build on prior HER2-focused mathematical models [27] by (i) explicitly capturing plastic interconversion between HER2+ and HER2-states under treatment, (ii) incorporating division-mode controls on heterogeneity [20], and (iii) exploring scheduling/sequencing of multi-agent regimens motivated by experimental synergy patterns [28] and eco-evolutionary competition principles [21]. We aim to generate testable predictions for combination designs that suppress recurrence by targeting both signaling (HER2/Notch) and state dynamics.

## 2 Model and Methods

### 2.1 Baseline two-phenotype model

We adopt the two-state framework of Li and Thirumalai [27], which tracks the populations of HER2-positive (HER2^+^) and HER2-negative (HER2^−^) tumor cells, denoted *N*_1_(*t*) and *N*_2_(*t*), with total size *N*(*t*) = *N*_1_(*t*) + *N*_2_(*t*). Each phenotype undergoes symmetric division at rates *K*_1_ (HER2^+^) and *K*_2_ (HER2^−^). Phenotypic plasticity is represented by asymmetric transitions at rates *K*_12_ (HER2^+^ →HER2^−^) and *K*_21_ (HER2^−^ →HER2^+^).

The asymmetric transition rates *K*_12_ and *K*_21_ are taken to be smaller than the symmetric division rates *K*_1_ and *K*_2_, reflecting the assumption that reversible phenotypic switching occurs less frequently than phenotype-preserving proliferation. This is consistent with recent experimental and modeling studies suggesting that such plastic transitions may occur only once every few cell divisions [29].

Here, symmetric division increases population size, while asymmetric transitions represent fate changes occurring during division that preserve the total cell number. In the absence of density regulation or therapy, the system is linear and grows exponentially, with long-time composition determined by the dominant eigenvector of the associated 2 × 2 matrix. Using the calibrated parameters of [27] (Table 1), trajectories converge robustly to a HER2^+^-dominant mixture of approximately (0.78, 0.22) across a wide range of initial conditions. This equilibrium composition is consistent with experimental observations in which sorted HER2^+^ and HER2^−^ populations reconstitute a stable mixed state over time. Importantly, this coexistence cannot be explained by differences in proliferation rates alone; rather, it arises from bidirectional phenotypic transitions between the two cell states, which are essential to maintain the observed equilibrium. Throughout, we assume: (i) no baseline apoptosis, and (ii) no symmetric events that produce two daughters of the opposite phenotype. Baseline apoptosis is omitted to isolate the effects of phenotypic plasticity and competitive growth; cell loss is introduced explicitly only through therapy in later sections.

**Table 1:**
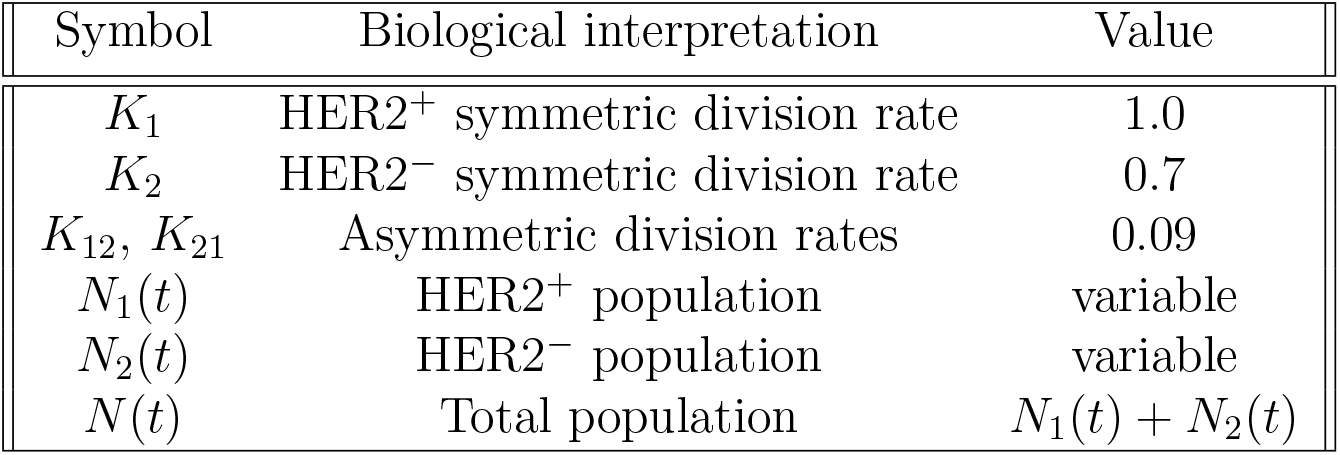
Baseline parameters for the Li–Thirumalai model (1).

| Symbol | Biological interpretation | Value |
| --- | --- | --- |
| $K_1$ | HER2<sup>+</sup> symmetric division rate | 1.0 |
| $K_2$ | HER2<sup>−</sup> symmetric division rate | 0.7 |
| $K_{12}, K_{21}$ | Asymmetric division rates | 0.09 |
| $N_1(t)$ | HER2<sup>+</sup> population | variable |
| $N_2(t)$ | HER2<sup>−</sup> population | variable |
| $N(t)$ | Total population | $N_1(t) + N_2(t)$ |

**Table 2:** Therapy regimens examined in the sequential and simultaneous treatment simulations. Paclitaxel doses (25 nM and 100 nM) and the targeted–therapy exposure level (25 *µ*g/mL) are chosen to reflect commonly used experimental ranges reported in the literature (e.g. [19, 28]). These regimens are used to explore the qualitative behaviour of the 4C ecological model under phenotype–specific treatment sequencing.

| Regimen | Chemotherapy (Paclitaxel dose) | Targeted therapy |
| --- | --- | --- |
| Chemo → Targeted | 25 nM | 25 $\mu\text{g}/\text{mL}$ |
| Chemo → Targeted | 100 nM | 25 $\mu\text{g}/\text{mL}$ |
| Targeted → Chemo | 25 nM | 25 $\mu\text{g}/\text{mL}$ |
| Targeted → Chemo | 100 nM | 25 $\mu\text{g}/\text{mL}$ |
| Simultaneous (Chemo + Targeted) | 25 nM | 25 $\mu\text{g}/\text{mL}$ |
| Simultaneous (Chemo + Targeted) | 100 nM | 25 $\mu\text{g}/\text{mL}$ |

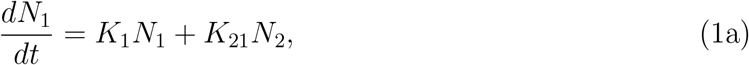

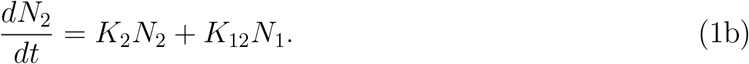

Closed-form expressions for eigenvalues, eigenvectors, and steady-state fractions appear in Appendix A.

### 2.2 Carrying-capacity extension: shared logistic regulation

To introduce density-dependent regulation while maintaining a minimal extension of the baseline dynamics, we first consider a symmetric logistic modification in which both phenotypes experience the same crowding factor 1 ^−^ *N*(*t*)*/C*, where *C* denotes a reference carrying-capacity scale. Rather than modeling contact inhibition, which is weak or absent in many cancer cells, this term is interpreted as a coarse limitation imposed by shared constraints such as available space, nutrients, oxygen, or vascular support within the tissue microenvironment.

The symmetric logistic extension is:

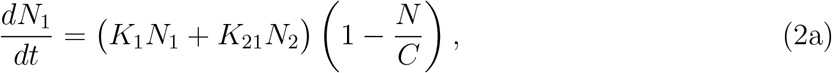

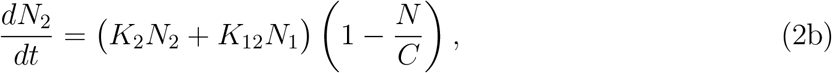

where *N*(*t*) = *N*_1_(*t*) + *N*_2_(*t*).

Because both phenotypes are regulated by the same crowding factor, density dependence acts only on the total population size and not on relative fitness. As a result, the phenotypic composition follows the same asymptotic proportions as in the linear Li–Thirumalai model, converging robustly to the mixture (0.78, 0.22), consistent with experimentally observed re-equilibration of HER2^+^ and HER2^−^ subpopulations driven by phenotypic plasticity, while the total population saturates at *C*. Because crowding acts identically on both phenotypes, this formulation cannot capture phenotype-specific competitive advantages. It therefore serves as a baseline and conceptual bridge to the more general competitive (4C) model introduced below.

### 2.3 Competition–coefficient extension (4C model)

The symmetric carrying-capacity model (2) cannot capture differential resource use or phenotype-specific ecological interactions. To incorporate these effects, we generalize the logistic term following Lotka–Volterra competition theory.

Let *C*_12_ denote the competitive load that HER2^−^ imposes on HER2^+^, and let *C*_21_ denote the reciprocal load. The resulting *4C model* is:

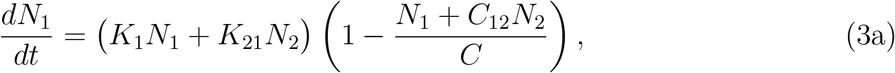

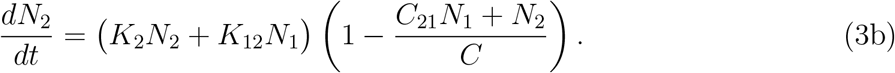

Here, *C* is a reference density scale that sets the magnitude of population size at which competitive interactions become dominant, rather than a hard carrying capacity. The *effective* carrying capacities differ across phenotypes, depending on *C*_12_ and *C*_21_, so the tumor’s saturation behavior depends on the competitive hierarchy rather than a fixed value of *C*.

When *C*_12_ = *C*_21_ = 1, the model reduces to the symmetric logistic system in Section 2.2. When 0 *< C*_12_, *C*_21_ *<* 1 (mutual weak competition), the system admits a biologically meaningful, stable coexistence equilibrium.

#### Early-time behaviour

For *N*(*t*) ≪ *C*, both crowding factors satisfy

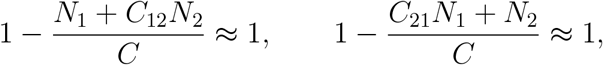

so the system behaves like the linear Li–Thirumalai model. Differences emerge only at high density, where competition alters the long-term phenotypic composition.

Throughout, we refer to the system as entering a “competition-dominated” or “saturated” regime when population size becomes comparable to the reference scale *C*, rather than implying a strict biological carrying capacity.

#### 2.3.1 4C Model Behavior

Assuming symmetric plasticity *K*_12_ = *K*_21_ = *K*_0_, the 4C system admits a coexistence equilibrium

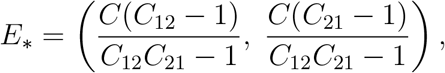

which is stable when 0 *< C*_12_, *C*_21_ *<* 1. Symmetric plasticity is assumed here to isolate the effects of the competition coefficients on equilibrium structure.

This equilibrium corresponds to a resource-limited coexistence state whose phenotypic composition is governed primarily by competitive asymmetry rather than intrinsic growth rates.

Depending on the competitive hierarchy:

- If *C*_21_ *> C*_12_, HER2^+^ dominates at high density.
- If *C*_12_ *> C*_21_, HER2^−^ dominates.
- If *C*_12_ = *C*_21_, a balanced coexistence arises.

This classification highlights a sharp sensitivity to the relative magnitude of *C*_12_ and *C*_21_. In particular, the symmetry line *C*_12_ = *C*_21_ acts as a threshold separating qualitatively distinct ecological regimes, so that even small deviations in competition strength can lead to a reversal in the dominant phenotype and, consequently, in therapeutic outcomes.

#### 2.3.2 Competition coefficient values

Guided by biological considerations and stability analysis, we restrict to the *weak-competition* regime

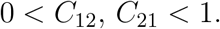

Here, *C*_12_ measures the inhibition of HER2^+^ by HER2^−^, and *C*_21_ the inhibition of HER2^−^ by HER2^+^. Weak cross-inhibition avoids competitive exclusion on experimental timescales.

Because HER2^+^ tumors are frequently linked to metabolic reprogramming and increased glucose and resource demand mediated by HER2–PI3K/AKT/mTOR signaling, it is biologically plausible to consider *C*_21_ ≳ *C*_12_ [30–32].

We do not attempt to infer precise competition coefficients; instead, we explore qualitative regimes consistent with weak mutual inhibition. The reverse ordering (*C*_12_ *> C*_21_) becomes important when examining long-term ecological dynamics or therapy initiated near saturation. In simulations, we consider representative weak-competition regimes using *C*_12_ = 0.4, *C*_21_ = 0.8 (HER2^+^-favoured), *C*_12_ = 0.8, *C*_21_ = 0.4 (HER2^−^-favoured), and *C*_12_ = *C*_21_ = 0.6 (symmetric competition). For simulations involving competition dynamics (Figs. 2, 4–7), we use the HER2^+^-favoured regime (*C*_12_, *C*_21_) = (0.4, 0.8) unless otherwise stated, while Fig. 8 explicitly compares all three regimes.

**Figure 1:**
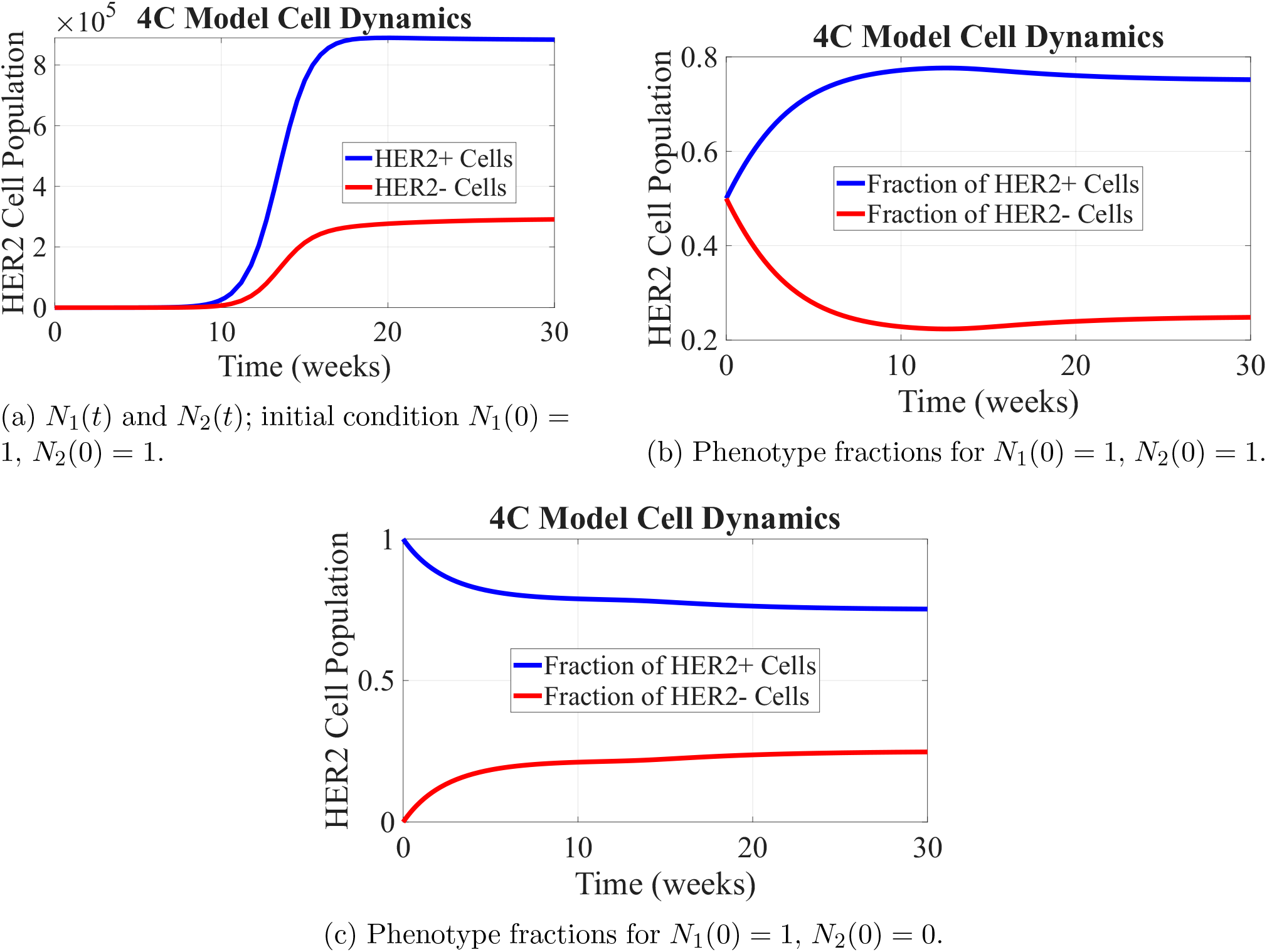
Dynamics under Model (2) with *C* = 10^6^. In all cases, the total population saturates due to shared resource limitation, while the phenotypic composition converges to the Li–Thirumalai mixture (0.78, 0.22), consistent with experimentally observed steady-state composition arising from bidirectional phenotypic plasticity.

**Figure 2:**
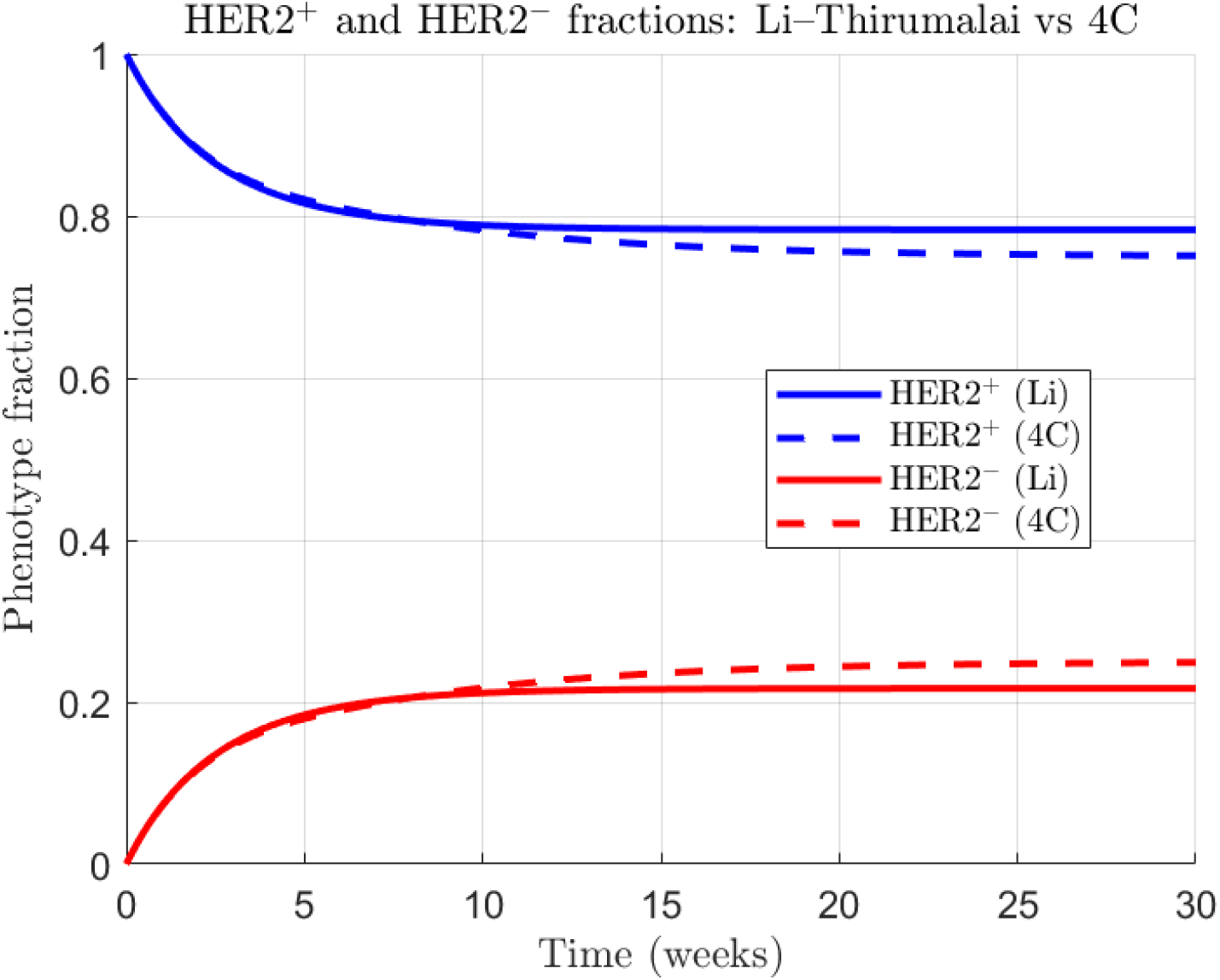
Comparison of phenotype fractions under the Li–Thirumalai model (1) (solid) and the 4C model (3) (dashed). Early-time trajectories coincide; at high density, asymmetric competition shifts the equilibrium composition. Simulations use (*C*_12_, *C*_21_) = (0.4, 0.8).

**Figure 3:**
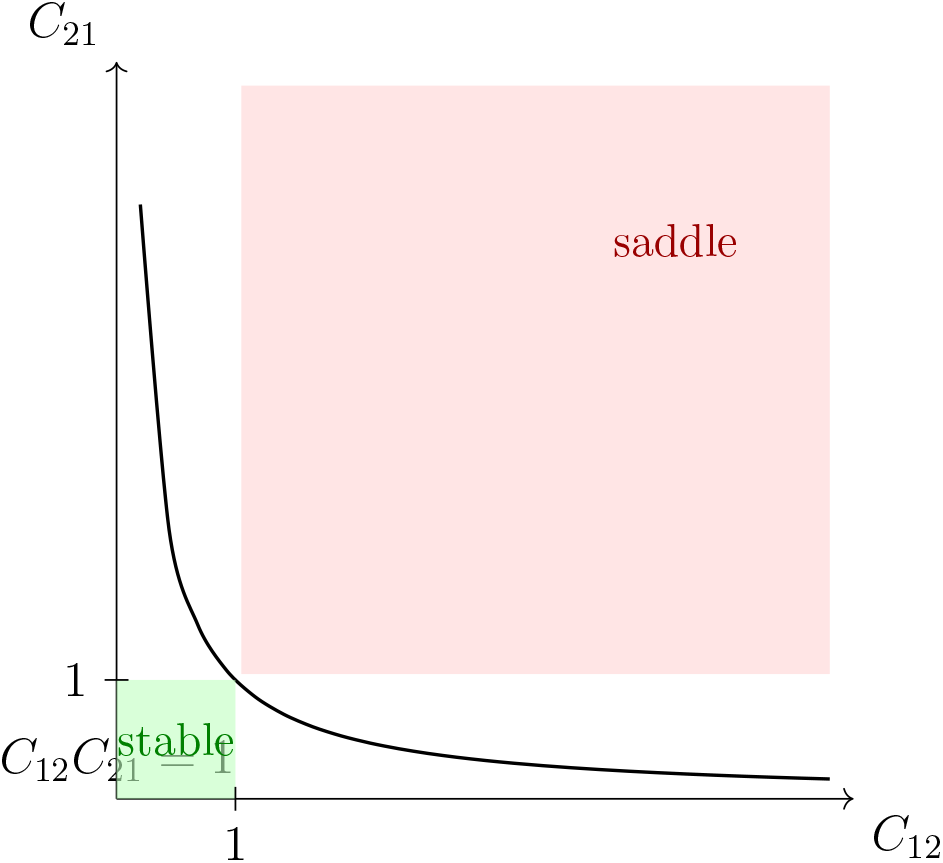
Stability map for the coexistence equilibrium of the 4C model in the (*C*_12_, *C*_21_) plane.

**Figure 4:**
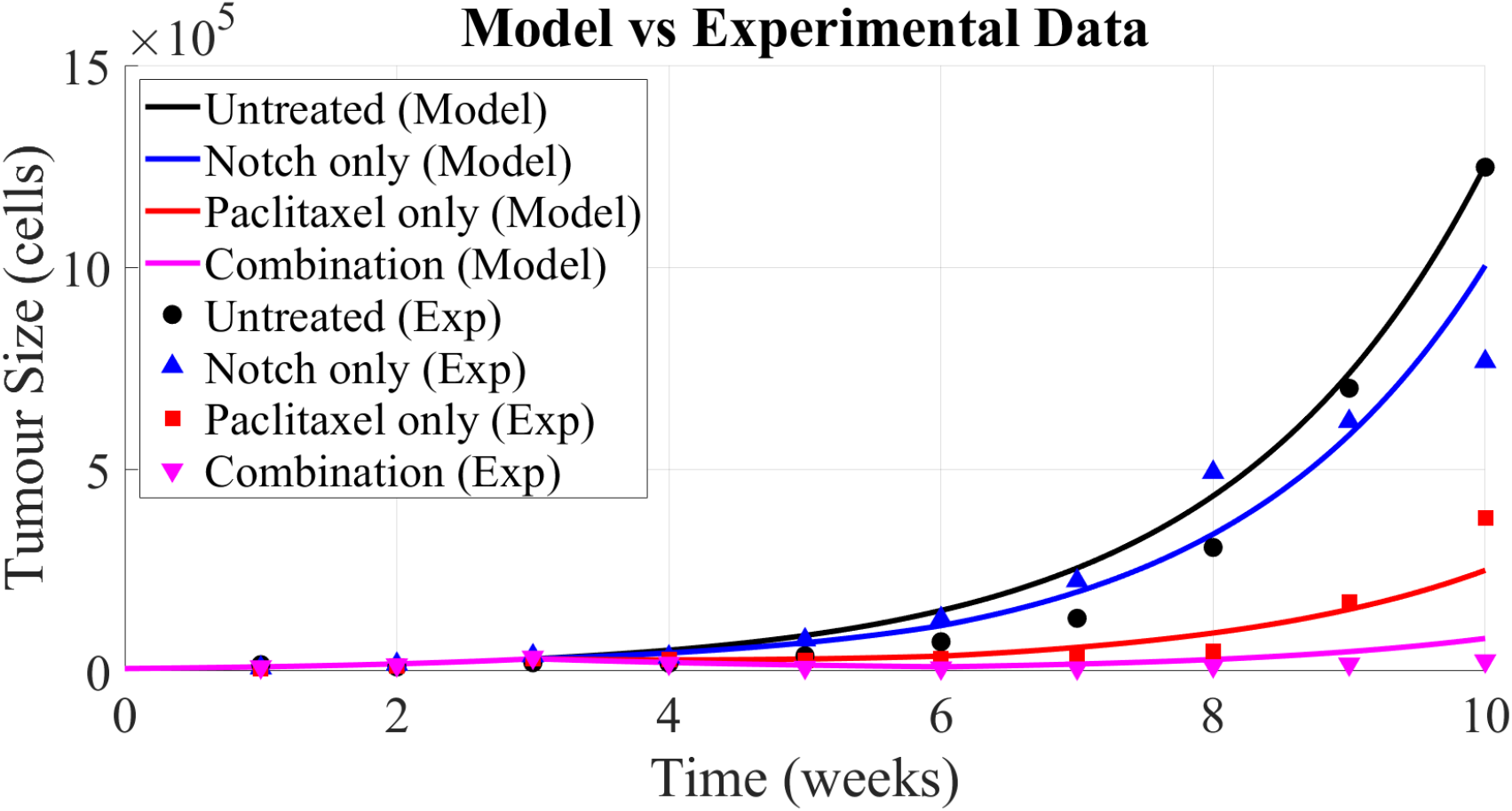
Comparison between model predictions and experimental tumour–growth data from Jordan *et al*. [19]. Solid lines denote model predictions for untreated growth (black), Notch–inhibitor monotherapy (blue), Paclitaxel monotherapy (red), and combined therapy (magenta). Symbols denote the corresponding experimental measurements, which were digitized from published figures using *WebPlotDigitizer* [33]. Treatments were applied during weeks 3–6 following the experimental protocol, with downstream effects persisting through the pharmacodynamic response. Simulations use initial fractions *N*_1_(0) : *N*_2_(0) = 0.78:0.22, corresponding to the experimentally observed steady-state composition arising from bidirectional phenotypic transitions, and in vivo growth rates rescaled following Li and Thirumalai [27]. The comparison demonstrates qualitative agreement between model predictions and experimental trends. Simulations use (*C*_12_, *C*_21_) = (0.4, 0.8).

### 2.4 Therapy Incorporation

Tumours composed of heterogeneous phenotypes are frequently treated using *combination therapy*, in which a cytotoxic agent targets rapidly proliferating cells while a targeted agent interferes with a phenotype–specific signalling pathway. Consistent with experimental observations in HER2–plastic breast cancer models [19, 27], we take Paclitaxel to act predominantly on HER2^+^ cells, while a Notch–pathway inhibitor primarily affects HER2^−^ cells.

In the original Li–Thirumalai formulation, therapeutic effects were encoded through a single effective parameter *γ*, which does not distinguish between drug mechanisms or phenotype specificity. Here, we instead integrate each therapy mechanistically into the nonlinear 4C ecological framework and then combine them. This enables phenotype–specific drug action while preserving nonlinear competition through the shared carrying capacity *C* and asymmetric interaction coefficients *C*_12_ and *C*_21_.

#### 2.4.1 Chemotherapy (Paclitaxel)

Paclitaxel is a microtubule–stabilizing cytotoxic agent that induces mitotic arrest and cell death, acting primarily on rapidly dividing HER2^+^ cells. Following prior pharmacodynamic formulations [28], we introduce a bounded pharmacodynamic (PD) variable *P*(*t*) that relaxes toward a prescribed dosing input *P*_*f*_(*t*). The parameter *τ*_*P*_ governs the rate at which intracellular drug activity responds to changes in external exposure. The relaxation time *τ*_*P*_ therefore captures intracellular drug response dynamics rather than systemic pharmacokinetics.

The HER2^+^ dynamics therefore acquire a drug–induced death term:

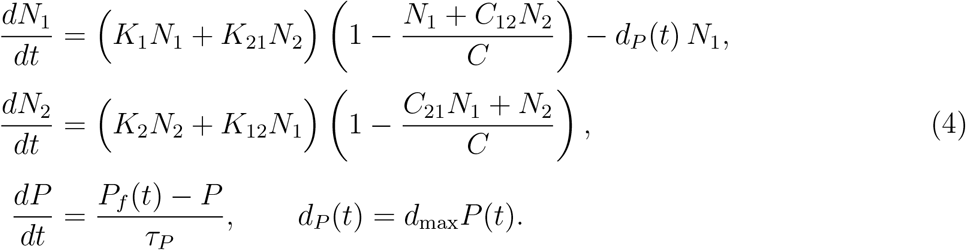

Here *d*_max_ denotes the maximal achievable HER2^+^ death rate under saturated Paclitaxel activity.

##### Dose–effect mapping

Experimental Paclitaxel concentrations are mapped to the normalized PD input *P*_*f*_(*t*) ∈ [0, 1] using a linear scaling,

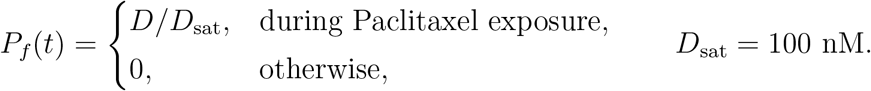

Thus 25 nM and 100 nM correspond to *P*_*f*_ = 0.25 and *P*_*f*_ = 1, respectively. The parameter *d*_max_ is calibrated so that the Paclitaxel–only simulations reproduce the experimentally observed HER2^+^ response under the 25 nM protocol [28].

#### 2.4.2 Targeted therapy (Notch inhibitor)

We next introduce a Notch–pathway inhibitor that acts selectively on HER2^−^ cells through receptor–mediated growth inhibition. Unlike standard inhibitory models that restrict suppression factors to the interval [0, 1], we explicitly allow the inhibition function *f*(*A*_*b*_, *t*) to exceed unity, representing net HER2^−^ population loss under sufficiently strong Notch signaling suppression. This formulation provides a compact phenomenological mechanism for capturing treatment–induced tumour regression without introducing a separate cytotoxic death term.

The pharmacodynamic structure is adapted from the binding–based model of Jarrett *et al*. [28], while the biological interpretation is guided by experimental studies demonstrating that strong Notch inhibition can induce transient tumour regression rather than mere growth delay [19].

The governing equations are

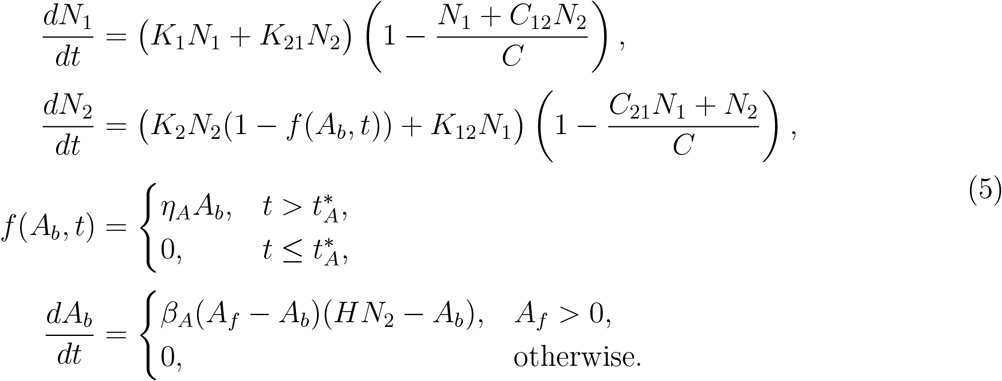

Here *A*_*b*_(*t*) denotes the amount of inhibitor bound to HER2^−^ receptors, *A*_*f*_(*t*) is the administered free drug, and *H* is the receptor abundance per HER2^−^ cell. The function *f*(*A*_*b*_, *t*) acts as an *effective inhibition factor* modulating the intrinsic HER2^−^ proliferation rate.

Allowing *f*(*A*_*b*_, *t*) *>* 1 corresponds to net negative growth of the HER2^−^ population under strong receptor occupancy and provides a biologically interpretable representation of treatment–induced cell loss. This extension is consistent with experimental observations of tumour regression under Notch inhibition and preserves the original binding–mediated structure of the Jarrett *et al*. model [19, 28].

In numerical simulations, the magnitude of *f*(*A*_*b*_, *t*) is controlled implicitly by the administered dose and receptor–binding dynamics, rather than by imposing an explicit upper bound or introducing a separate cytotoxic death term.

#### 2.4.3 Combination therapy

We finally combine both therapeutic mechanisms within the 4C framework. The two therapies act on distinct phenotypes through independent pharmacodynamic variables: Paclitaxel induces cytotoxic death of HER2^+^ cells, while the Notch inhibitor suppresses HER2^−^ proliferation and, under sufficiently strong inhibition, can induce net negative growth.

The resulting coupled system is

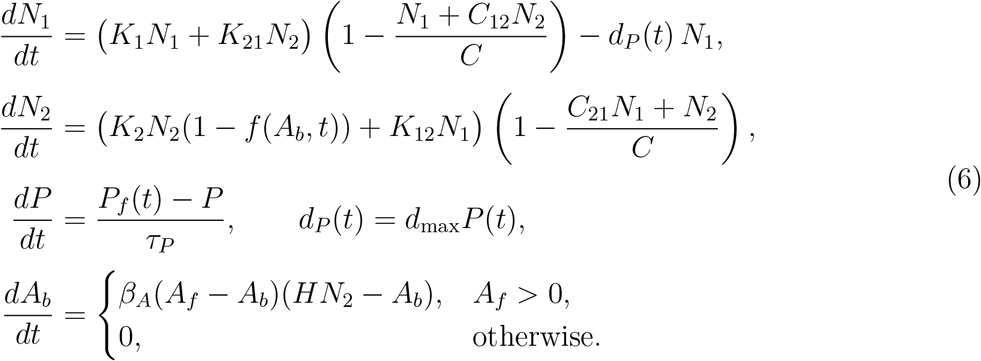

In the absence of therapy (*P*_*f*_ ≡ *A*_*f*_ ≡ 0), both pharmacodynamic variables relax to zero and the system reduces exactly to the untreated 4C model. This modular extension preserves the validated baseline dynamics of the Li–Thirumalai framework while enabling phenotype–specific treatment responses consistent with experimental observations.

All simulations were implemented in MATLAB; representative code supporting this study is publicly available at [GitHub](https://github.com/GAVRI015/tumor-heterogeneity-paper-code).

## 3 Results

We first evaluated the ability of the revised therapy–integrated 4C model to reproduce the experimentally observed tumour–growth dynamics reported by Jordan *et al*. [19] under four treatment conditions: untreated, Notch–inhibitor monotherapy, Paclitaxel monotherapy, and the simultaneous combination of both agents. For each condition, we solved the coupled population–pharmacodynamic system using treatment schedules matching the experimental protocol (weeks 3–6) and compared the predicted total tumour–cell number with the bioluminescent photon–flux measurements (Fig. 4).

To initialise the xenografts, we set the HER2^+^/HER2^−^ composition to the experimentally supported equilibrium ratio 0.78:0.22, which arises from bidirectional phenotypic transitions between the two cell states and is captured by the dominant eigenstructure of the Li–Thirumalai linear system. The initial total tumour burden was chosen so that the untreated simulation matches the first–week photon–flux measurement. Because absolute tumour cell numbers are not directly reported in the xenograft experiments, we fixed the initial total population at *N*(0) = 5,000 cells, which provides a consistent scale for both the experimental data and the population sizes used in the Li–Thirumalai framework. Importantly, model predictions depend on relative population fractions and competitive interactions rather than absolute cell counts.

Following [27], we rescaled the intrinsic symmetric division rates according to

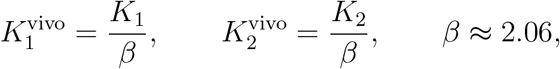

which aligns the untreated in vivo growth rate with the xenograft data. Importantly, the resulting untreated baseline trajectory—including the slow approach toward a HER2^+^–dominant composition—is consistent with the short–time behaviour reported by Li and Thirumalai [27], confirming that the 4C framework preserves validated tumour dynamics prior to the introduction of therapy.

Across all treatment conditions, the model captures the key qualitative and quantitative features of the experimental responses. In the absence of treatment, the total tumour burden exhibits sustained exponential–like expansion, consistent with the observed growth of xenografts derived from mixed HER2^+^/HER2^−^ circulating tumour cells. Under Paclitaxel monotherapy, the bounded pharmacodynamic surrogate acting selectively on HER2^+^ cells induces an early decline in tumour burden, followed by regrowth once the drug effect relaxes, closely matching the experimental trend.

Notch–inhibitor monotherapy produces a more moderate but sustained suppression of tumour growth. In the model, this response arises from receptor–mediated inhibition of HER2^−^ proliferation, which under sufficiently strong exposure can transiently induce net negative growth of the HER2^−^ population. This mechanism enables partial tumour regression during treatment while preserving the capacity for regrowth following drug withdrawal, consistent with the experimental observations.

The combined therapy produces the strongest and most durable tumour suppression. The model successfully reproduces both the depth and duration of tumour reduction observed experimentally, reflecting the complementary actions of Paclitaxel–induced HER2^+^ cell death and strong Notch–mediated suppression of HER2^−^ dynamics. From an ecological perspective, this synergy arises from simultaneously suppressing both phenotypes before competitive release can occur, thereby preventing the untreated subpopulation from exploiting therapy–induced niche availability.

Overall, these results demonstrate that the revised 4C model—integrating distinct pharmacodynamic mechanisms for chemotherapy and targeted therapy—can quantitatively reproduce the heterogeneous treatment responses observed *in vivo*. The framework provides a coherent and biologically interpretable platform for analysing phenotype–specific therapeutic interactions and for assessing how treatment strength and scheduling shape tumour evolution in heterogeneous cancers.

### 3.1 Sequential and simultaneous therapy schedules

We next examined how the revised therapy–integrated 4C model responds to different orders of drug administration. Motivated by experimentally relevant treatment protocols reported in the literature, we considered three clinically meaningful schedules: (i) chemotherapy followed by targeted therapy, (ii) targeted therapy followed by chemotherapy, and (iii) simultaneous administration of both agents. While the xenograft measurements analysed here are drawn from Jordan *et al*. [19], the dose ranges and exposure windows explored are informed by commonly used experimental protocols, including those reported by Jarrett *et al*. [28].

In our framework, each treatment is implemented as a pharmacodynamic pulse through the variables *P*(*t*) (Paclitaxel activity) and *A*_*b*_(*t*) (bound Notch inhibitor). This abstraction allows us to focus on the population–level consequences of treatment ordering and phenotype–specific drug action, rather than reproducing detailed pharmacokinetic timescales.

Although experimental studies tested multiple dose combinations, our aim here is not to quantitatively refit xenograft growth curves, but to explore the qualitative behaviour of the 4C ecological model under representative chemotherapy and targeted–therapy intensities. Accordingly, we examine two Paclitaxel exposure levels (25 nM and 100 nM), each paired with a single targeted–therapy level of 25 *µ*g/mL.

Simulations are initialised at the HER2^+^/HER2^−^ equilibrium composition *N*_1_(0) : *N*_2_(0) = 0.78 : 0.22 predicted by the Li–Thirumalai eigenstructure and are run over a 20–week horizon, with all therapy confined to weeks 3–7. Because treatment is applied well below saturation during this early interval, the total tumour burden remains far from the competition–dominated regime. Ecological interactions are therefore latent rather than dominant, and short–term dynamics are driven primarily by phenotype–specific drug action. Density–dependent competition becomes relevant only at later times, shaping long–term regrowth and phenotypic composition after therapy cessation.

#### 3.1.1 Chemotherapy prior to targeted therapy

In the chemo → targeted sequence, Paclitaxel produces a rapid decline in the HER2^+^ subpopulation, reflecting its selective cytotoxic action (Figs. 5a–5b). Once the chemotherapy pulse ends, HER2^+^ cells recover through symmetric division and phenotypic interconversion, leading to renewed tumour growth that remains well below the competition–dominated regime.

**Figure 5:**
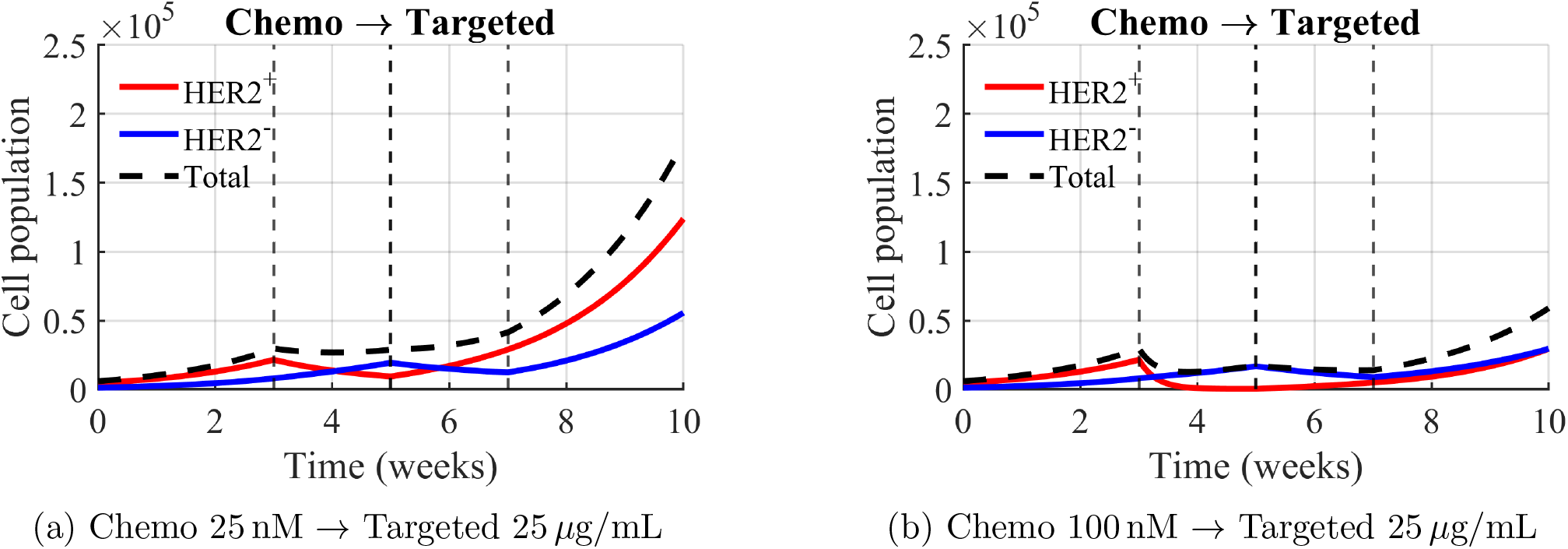
Sequential chemotherapy→targeted therapy schedules. Paclitaxel is administered from weeks 3–5, followed by targeted therapy from weeks 5–7. Simulations use (*C*_12_, *C*_21_) = (0.4, 0.8).

**Figure 6:**
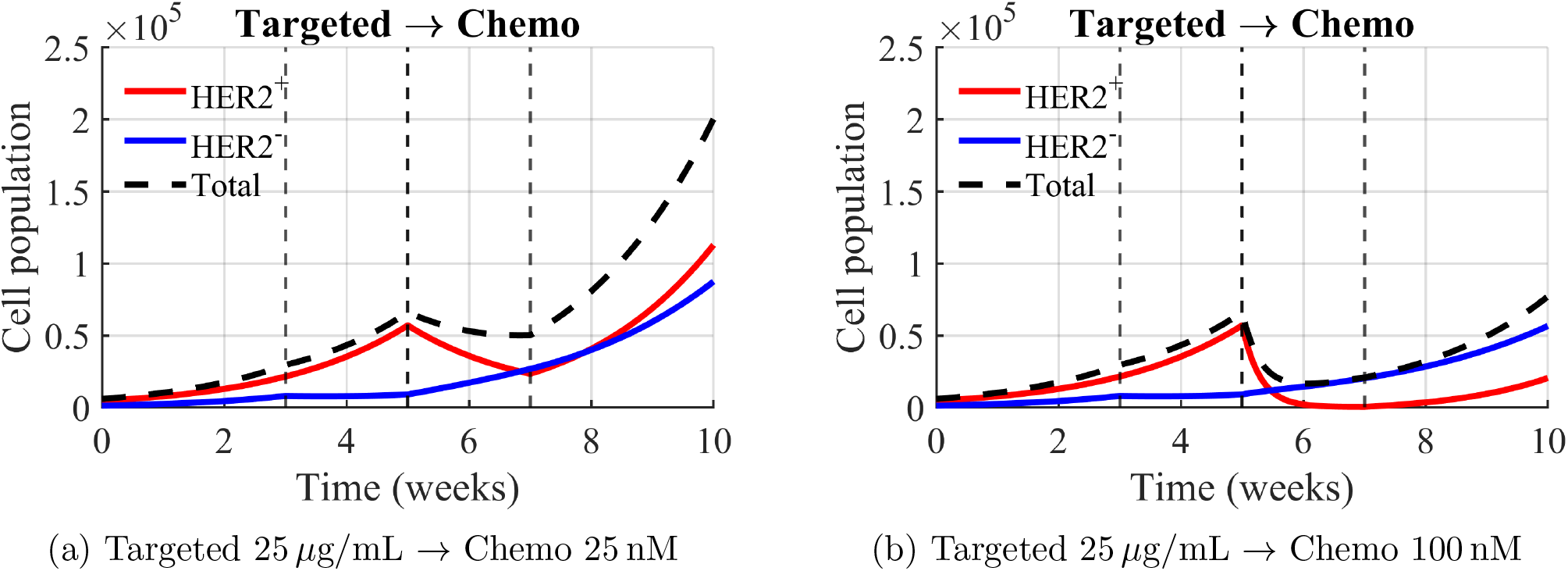
Sequential targeted→chemotherapy schedules. Targeted therapy is applied during weeks 3–5, followed by Paclitaxel from weeks 5–7. Simulations use (*C*_12_, *C*_21_) = (0.4, 0.8).

Introduction of targeted therapy at week 5 suppresses HER2^−^ proliferation, slowing subsequent expansion. Increasing the Paclitaxel dose amplifies the initial HER2^+^ depletion and delays post–treatment regrowth, but the qualitative behaviour is preserved across doses.

#### 3.1.2 Targeted therapy prior to chemotherapy

When targeted therapy is administered first, HER2^−^ proliferation is suppressed during the initial treatment window, producing a moderate reduction in tumour growth. Subsequent Paclitaxel administration sharply depletes the HER2^+^ population, driving a pronounced reduction in total tumour burden.

Higher Paclitaxel doses deepen HER2^+^ depletion and delay recovery, but in all cases short–term dynamics are governed by phenotype–specific drug action, with competitive effects remaining weak until later stages of regrowth.

#### 3.1.3 Simultaneous therapy

Figure 7 shows that simultaneous delivery of targeted therapy and Paclitaxel produces coordinated suppression of both HER2^+^ and HER2^−^ populations throughout the treatment window. Total tumour burden peaks near treatment onset and then declines rapidly once both agents are applied.

**Figure 7:**
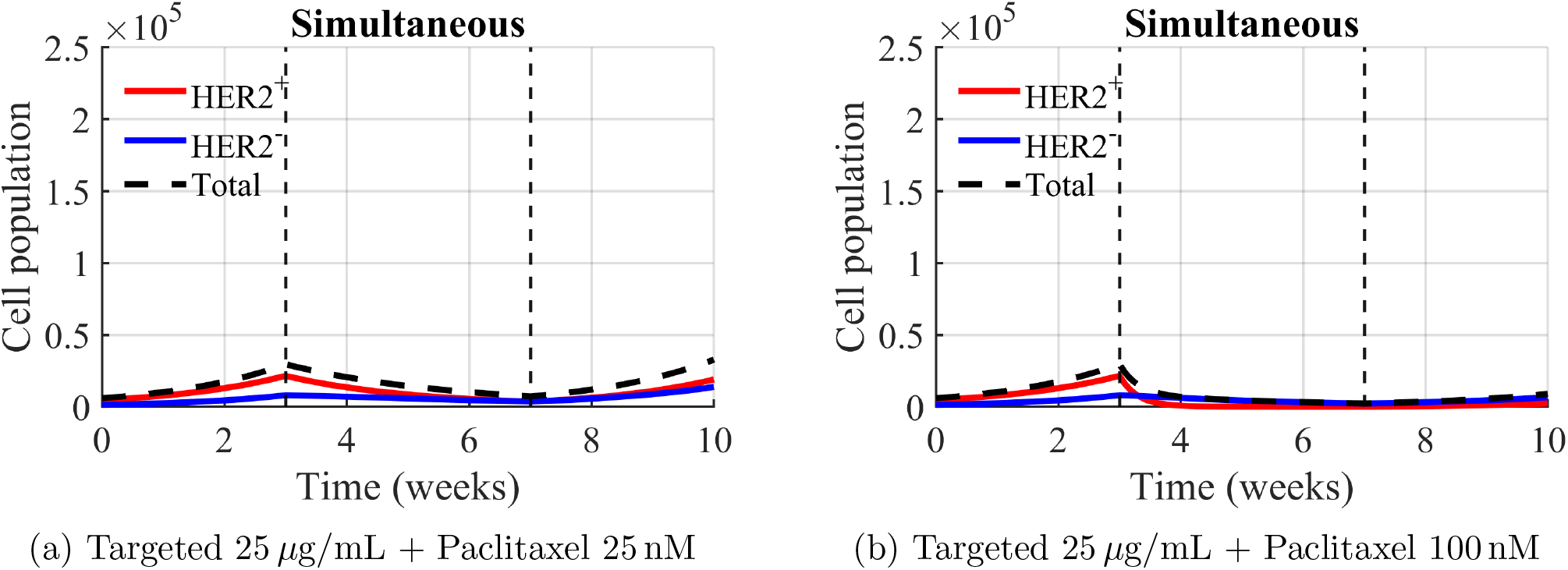
Simultaneous co-treatment from weeks 3–7. Simulations use (*C*_12_, *C*_21_) = (0.4, 0.8).

**Figure 8:**
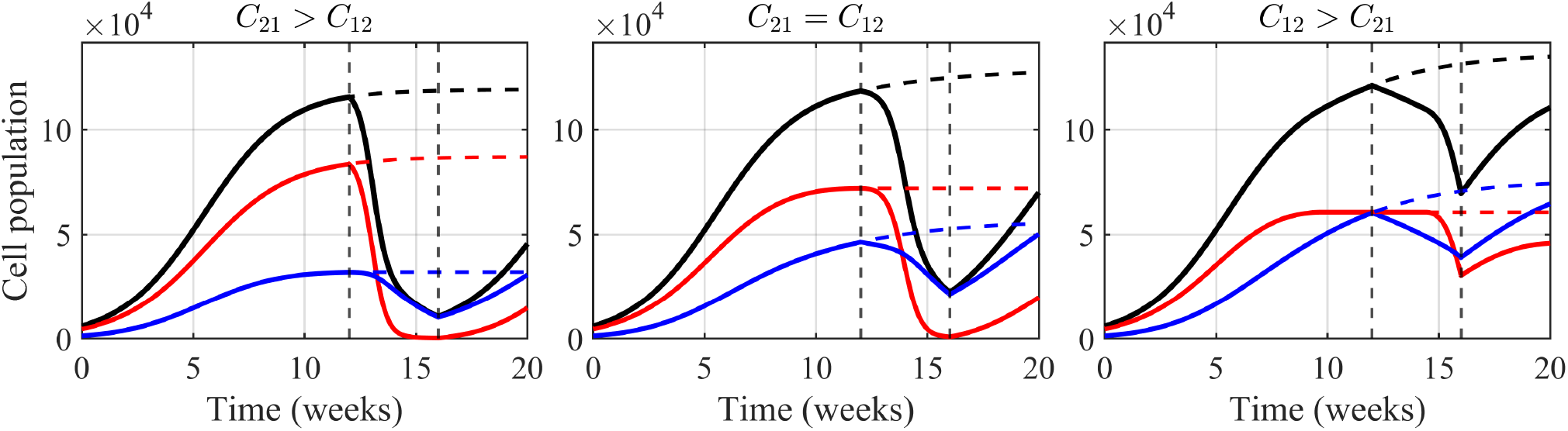
Treatment initiated in the competition-dominated regime. Solid curves show treated dynamics for HER2^+^(red), HER2^−^(blue), and total tumour burden (black); dashed curves indicate the corresponding untreated trajectories. Simultaneous Paclitaxel and Notch inhibition is applied from weeks 12–16 (vertical dashed lines). Panels correspond to different competitive hierarchies: *C*_21_ *> C*_12_(left, (*C*_12_, *C*_21_) = (0.4, 0.8)), *C*_21_ = *C*_12_(middle, (*C*_12_, *C*_21_) = (0.6, 0.6)), and *C*_12_ *> C*_21_(right, (*C*_12_, *C*_21_) = (0.8, 0.4)).

Under the lower Paclitaxel dose, both phenotypes persist at reduced levels, leading to gradual post–therapy regrowth. At the higher dose, HER2^+^ cells are nearly eradicated and tumour burden approaches zero by week 7, producing deeper and more durable suppression.

Because the tumour remains well below the competition–dominated regime during and shortly after treatment, ecological interactions do not strongly influence these early dynamics. Instead, post–therapy behaviour is governed primarily by intrinsic regrowth rates, with HER2^−^ cells recovering earlier and HER2^+^ cells eventually regaining dominance due to their higher proliferation capacity. As a result, the system does not immediately return to a competition-set equilibrium; such behaviour would only emerge at later times if the tumour regrows to sufficiently high density.

### 3.2 Treatment initiated in the competition-dominated regime

Whereas early-treatment dynamics are governed primarily by drug specificity and phenotypic plasticity, late-treatment dynamics are shaped by ecological constraints imposed by inter-phenotype competition. To assess how these constraints influence therapeutic efficacy, we examined treatment schedules initiated only after the tumour had entered a high-density regime in which crowding substantially limits net expansion.

In this setting, phenotypic transitions remain active, but the relative magnitudes of the competition coefficients *C*_12_ and *C*_21_ determine which phenotype is ecologically favoured. As a result, identical treatment schedules can produce qualitatively distinct outcomes depending solely on the underlying competitive hierarchy.

Accordingly, we considered three biologically plausible competition regimes: (i) *C*_21_ *> C*_12_, favouring HER2^+^ dominance, (ii) *C*_21_ = *C*_12_, corresponding to neutral coexistence, and (iii) *C*_12_ *> C*_21_, favouring HER2^−^ dominance.

For each regime, the tumour was first simulated without treatment until the dynamics approached a competition-dominated plateau characterised by marked slowing of total population growth and stabilisation of phenotype fractions. From this high-density state, we then applied *simultaneous* Paclitaxel and Notch inhibition during weeks 12–16 and compared the treated dynamics with the corresponding untreated trajectories (Fig. 8).

In the HER2^+^-favoured regime (*C*_21_ *> C*_12_; Fig. 8, left), chemotherapy induces a sharp transient decline in the HER2^+^ population and, consequently, in total tumour burden. However, suppression of the dominant phenotype relaxes competitive pressure on HER2^−^ cells, allowing their expansion during the treatment window. Following treatment cessation, phenotypic plasticity and competitive release restore a HER2^+^-dominant composition, yielding rapid regrowth. This behaviour illustrates *ecological resistance*.

In the neutral regime (*C*_21_ = *C*_12_; Fig. 8, middle), both phenotypes experience comparable competitive constraints. Treatment produces a pronounced but temporary reduction in tumour burden, after which the system relaxes back toward coexistence. This outcome reflects *conditional sensitivity*, where therapy alters short-term dynamics without overcoming ecological balance.

In the HER2^−^-favoured regime (*C*_12_ *> C*_21_; Fig. 8, right), the untreated tumour is dominated by HER2^−^ cells under crowding. Although Notch inhibition weakens the dominant phenotype during treatment, the reduction in total tumour burden is modest and recovery is rapid. HER2^+^ cells are not eradicated, demonstrating that late-applied therapy can yield only limited and transient reduction when the competitive hierarchy remains intact. We emphasize that this reduced effectiveness does not imply that the Paclitaxel–Notch inhibitor combination is intrinsically ineffective in HER2^−^–dominant tumours. Rather, this behaviour arises specifically when therapy is applied in a competition-dominated (high-density) regime in which the ecological hierarchy favours HER2^−^ cells. In this setting, competitive interactions and ongoing phenotypic plasticity limit tumour reduction and promote rapid recovery, reflecting ecological constraints imposed by competition rather than failure of the drug mechanism itself.

Taken together, these results demonstrate that once tumours are crowded, *competitive hierarchy* becomes a decisive determinant of late-time therapeutic response. Identical treatment schedules can yield transient debulking with rebound, moderate suppression with coexistence recovery, or weak suppression depending solely on the relative strengths of cross-competition. To our knowledge, this is among the first modelling frameworks to show that competitive hierarchy alone can qualitatively reshape the outcome of late-applied combination therapy.

## 4 Discussion

This study demonstrates that therapeutic response in a heterogeneous HER2^+^/HER2^−^ tumour is shaped not only by drug action, but by the interaction between phenotype-specific therapy mechanisms, ongoing phenotypic plasticity, and ecological competition within the 4C framework. A central insight is the clear separation between an early, therapy-dominated regime—where short-term dynamics are governed primarily by treatment selectivity—and a late, competition-dominated regime—where ecological constraints largely determine long-term outcome.

When treatment is initiated while the tumour remains well below carrying capacity, population dynamics are driven mainly by the direct effects of chemotherapy and targeted therapy on their respective phenotypes. In this early regime, different treatment orders and dose levels can produce similar short-term responses, reflecting the bounded and phenotype-selective nature of the pharmacodynamic effects. However, even when one phenotype appears strongly suppressed, neither population is permanently eliminated. Phenotypic plasticity maintains a low-level reservoir that allows regrowth once treatment pressure is relaxed, explaining why early apparent clearance does not translate into durable control.

As treatment continues, differences in ordering and relative intensity become important because they reshape competitive interactions between phenotypes. Aggressive chemotherapy applied after targeted therapy can reduce the competitive pressure imposed by HER2^+^ cells, unintentionally favouring HER2^−^ expansion through competitive release combined with ongoing phenotype switching. Conversely, more balanced dosing or simultaneous initiation can mitigate this effect by acting on both phenotypes before either gains a competitive advantage. Within the 4C framework, these non-monotone responses arise naturally from competition and plasticity, without invoking intrinsic drug resistance.

Simultaneous administration of chemotherapy and targeted therapy tends to produce stronger short-term suppression of total tumour burden than staggered schedules, as both phenotypes are acted upon concurrently. Nevertheless, phenotypic transitions remain active, so long-term outcome still depends on how competition reshapes the system once treatment pressure is removed. These findings suggest a practical strategy in which an initial combined pulse reduces both subpopulations, followed by maintenance therapy designed to prevent the ecologically favoured phenotype from exploiting competitive release.

The strongest evidence for ecological control of treatment outcome emerges when therapy is applied after the tumour has entered a crowded, competition-dominated state. As illustrated in Fig. 8, identical late-applied combination therapy can produce qualitatively distinct responses depending solely on the competitive hierarchy. When *C*_21_ *> C*_12_, transient suppression of HER2^+^ relaxes competitive pressure on HER2^−^, enabling compensatory growth during the treatment window and rapid rebound after cessation. When *C*_21_ = *C*_12_, treatment yields temporary suppression followed by recovery toward coexistence. When *C*_12_ *> C*_21_, HER2^−^ retains ecological dominance and overall tumour reduction is modest, with rapid post-treatment recovery. These outcomes show that late therapy can fail not because drug action is weak, but because ecological structure channels the system back toward its competition-set composition. Taken together, these results highlight two general principles. First, treatment order and relative intensity matter because they determine which phenotype experiences competitive release and when. Second, once tumours are crowded, competitive hierarchy becomes a decisive determinant of therapeutic response, independent of intrinsic proliferation rates. Importantly, this qualitative reversal arises without invoking acquired resistance, spatial heterogeneity, or time-varying drug sensitivity, and is driven solely by density-dependent competition interacting with ongoing phenotypic plasticity.

The therapy representations considered here are intentionally compact phenomenological surrogates that preserve biological interpretability while capturing experimentally observed trends. Future work will focus on formal dose–schedule optimisation within the 4C framework, using control-theoretic or Bayesian approaches to quantify trade-offs between tumour suppression and ecological rebound, and on assessing robustness across parameter uncertainty and patient-specific heterogeneity.

## 5 Conclusions

We developed a compact, mechanistically grounded ODE framework that preserves the phenotypic plasticity kinetics of the Li–Thirumalai model while embedding density-limited growth through a common carrying capacity and its asymmetric Lotka–Volterra generalization (the 4C model). Chemotherapy (Paclitaxel) and targeted Notch-pathway inhibition are incorporated as distinct, phenotype-specific actions via simple pharmacodynamic surrogates. Within this unified framework, we examined clinically relevant treatment schedules—chemo targeted, targeted chemo, and simultaneous dosing—and demonstrated that both treatment order and relative dose intensity play a decisive role in shaping long-term tumour composition and burden.

### Key findings

Four consistent observations emerge. First, treatment order matters: while short-term responses can appear similar across schedules and doses, long-term outcomes di-verge substantially once ecological feedback becomes important. Second, chemo-first schedules most effectively suppress the HER2^+^ compartment, particularly at higher chemotherapy doses, but leave the HER2^−^ population largely intact. Third, targeted-first schedules can exhibit *competitive release*, whereby stronger subsequent chemotherapy unintentionally favours HER2^−^ expansion through reduced cross-competition combined with ongoing phenotype switching. Fourth, simultaneous initiation largely mirrors the chemo-first response in suppressing HER2^+^, but with slower equilibration and without the pronounced HER2^−^ rebound seen in some targeted→chemo sequences. Taken together, these findings support a clear design principle: *begin with simultaneous chemotherapy and targeted therapy to curb both phenotypes, then maintain control with continued targeted inhibition*.

### Biological interpretation

The stability structure of the 4C model favours regimes of weak mutual competition (0 *< C*_12_, *C*_21_ *<* 1), consistent with resource-limited growth in which cross-inhibition is weaker than self-crowding. Within this ecologically plausible regime, sequence-dependent treatment outcomes arise naturally from the interaction of differential drug sensitivities, competitive dynamics, and bidirectional phenotypic plasticity. The model therefore provides a mechanistic explanation for why intensifying therapy does not necessarily improve control and can, in some cases, promote dominance of the less-sensitive phenotype.

### Limitations

The present analysis focuses on well-mixed, single-site tumour dynamics with a shared carrying capacity, symmetric plasticity calibration, single-dose treatment surrogates, and no explicit representation of toxicity or host constraints. These simplifications may underrepresent spatial heterogeneity, microenvironmental structure, drug–drug interactions, and resistance mechanisms beyond phenotypic plasticity. As such, quantitative predictions should be interpreted as mechanism-revealing rather than patient-specific.

In addition, phenotypic plasticity is represented through constant baseline transition rates *K*_12_ and *K*_21_, while treatment acts only through phenotype-specific pharmacodynamic effects. Thus, the model does not explicitly capture drug-induced changes in cell-state transition rates or the emergence of additional therapy-associated phenotypes. Recent experimental and theoretical studies have shown that anticancer therapies can alter phenotypic transition dynamics and promote drug-tolerant or hybrid cell states [34–36]. Incorporating treatment-dependent plasticity and additional emergent phenotypes is an important direction for future work.

Furthermore, the competition coefficients *C*_12_ and *C*_21_ are treated as fixed parameters, whereas therapy may alter inter-phenotype competition through changes in resource usage or microenvironmental interactions. Allowing these coefficients to vary dynamically under treatment could introduce additional ecological feedback and modify the competition-driven behaviours identified here.

Finally, the effects of Paclitaxel and Notch inhibition are modeled independently, without explicitly incorporating synergistic or antagonistic drug interactions. While this allows us to isolate the roles of phenotypic plasticity and ecological competition, incorporating drug–drug interactions may further influence treatment dynamics and represents an important direction for future work.

### Outlook

The 4C framework is readily extensible. Natural next steps include spatial reaction–diffusion or agent-based hybrids to capture niche structure and gradients; data-driven or time-varying plasticity laws; multi-dose PK/PD representations; and optimal-control or Bayesian approaches to personalise treatment intensity and scheduling under uncertainty. Prospective *in vitro* and *in vivo* studies—specifically probing the predicted advantages of simultaneous initiation with targeted maintenance and quantifying competitive release windows—will be essential for refining parameters and testing the robustness of these ecology-informed scheduling strategies.

## A Eigenstructure of the 2-by-2 Rate Matrix and Long–Run Composition

### Setup

For the baseline two-phenotype model (1), write

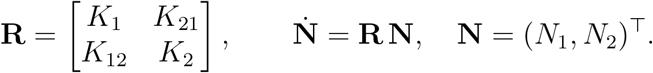

### Characteristic polynomial and closed-form eigenvalues

Eigenvalues solve det(**R** − *λI*) = 0, i.e.

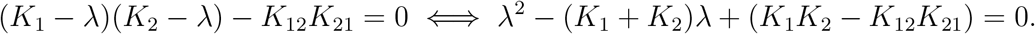

Hence

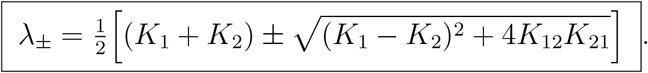

### Dominant right eigenvector

Let *λ*_+_ be the larger (dominant) eigenvalue. Any right eigenvector *v*_+_ ≠ 0 satisfies (**R** − *λ*_+_*I*)*v*_+_ = 0. Two convenient equivalent forms are

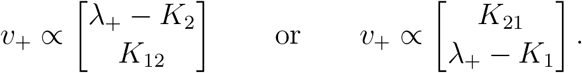

### Long–run fractions (composition)

Normalize *v*_+_ so its components sum to one:

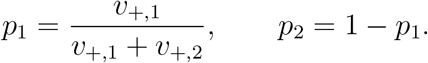

Using the first eigenvector form gives the closed formulas

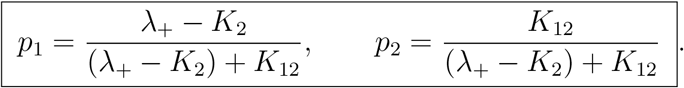

(Using the alternate form yields the equivalent 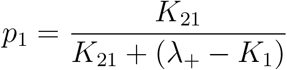)

### Numerical check (baseline calibration)

With *K*_1_ = 1.0, *K*_2_ = 0.7, *K*_12_ = *K*_21_ = 0.09, *λ*_+_ ≈ 1.02493, *p*_1_ ≈ 0.783, *p*_2_ ≈ 0.217, matching the ∼ (0.78, 0.22) HER2^+^/HER2^−^ mixture cited in the main text.

### Stability interpretation (linear system)

For 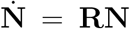: the extinction state **N** = **0** is asymptotically stable iff both eigenvalues have negative real parts. Here *λ*_±_ *>* 0, so **0** is *unstable*; however, *directions* converge to *v*_+_(Perron–Frobenius), yielding the stable fractions (*p*_1_, *p*_2_).

### Shared logistic factor

Multiplying the RHS by a common crowding term 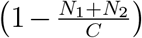 does not change eigenvectors; thus the limiting composition on the carrying-capacity manifold is still (*p*_1_, *p*_2_).

### Connection to the 4C model

With asymmetric competition coefficients (Eqs. (3)), the line of equilibria collapses to a single coexistence equilibrium *E*_*_. For 0 *< C*_12_, *C*_21_ *<* 1, *E*_*_ is locally asymptotically stable; for *C*_12_, *C*_21_ *>* 1 it is a saddle.

## B 4C model steady-state analysis (details)

### Jacobian

For the 4C system with *K*_0_ = *K*_12_ = *K*_21_, the Jacobian at (*N*_1_, *N*_2_) is

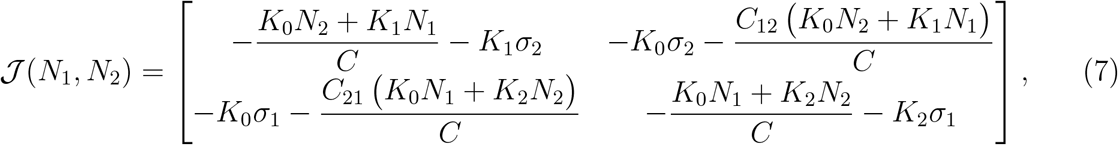

where

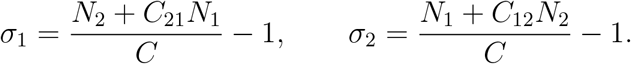

### Equilibria

Solving *dN*_1_*/dt* = *dN*_2_*/dt* = 0 yields

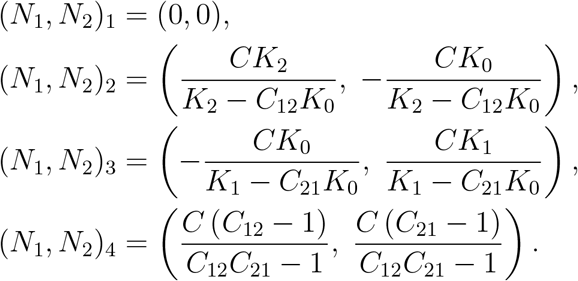

Under the Li–Thirumalai baseline, (*N*_1_, *N*_2_)_2_ and (*N*_1_, *N*_2_)_3_ are not biologically feasible.

### Eigenvalues at the extinction state

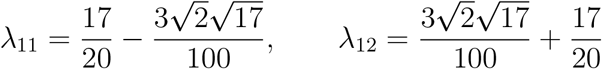

so that numerically *λ*_11_ ≈ 0.675, *λ*_12_ ≈ 1.025 (unstable).

### Eigenvalues at the coexistence state

A representative closed form is

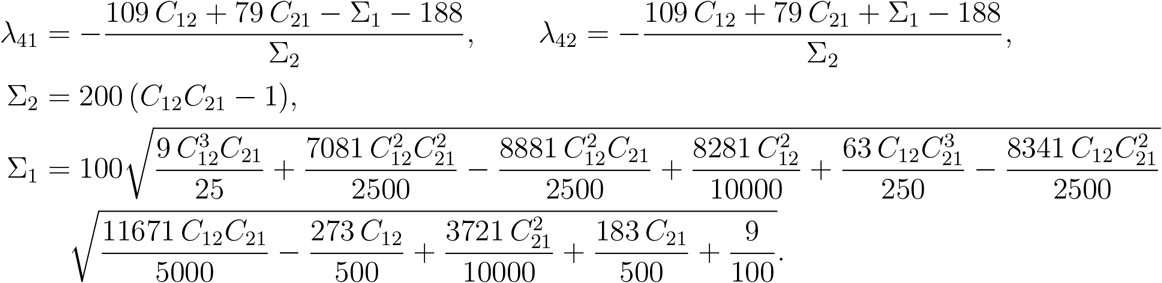

For figures and sign tests we evaluate these numerically in MATLAB (code provided in the supplement); no additional reparameterized numeric form is needed here.

### Stability classification

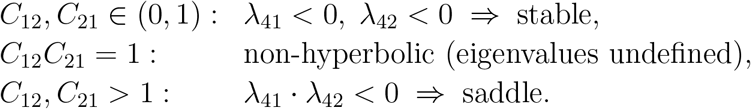

## Notes

https://github.com/GAVRI015/tumor-heterogeneity-paper-code

### Competing Interest Statement

The authors have declared no competing interest.

### Summary of Updates

This revised version improves the clarity and organization of the manuscript. The modeling framework and its biological motivation have been further clarified, and the presentation of results has been refined. Figure captions have been revised to provide clearer explanations and improve readability, while the figures themselves remain unchanged. A code availability statement has been added, and the simulation code is now publicly available to support reproducibility.

## References

[1] American Cancer Society, “Cancer facts & figures 2025,” Atlanta, GA, 2025, american Cancer Society.

[2] National Cancer Institute, “SEER cancer stat facts: Common cancer sites,” National Institutes of Health, Jul. 2025, accessed: 2025-07-15.

[3] M. S. Shiels et al., “Trends in cancer incidence and mortality rates in early-onset and older-onset age groups in the united states, 2010–2019,” Cancer Discovery, vol. 15, no. 5, pp. 1123–1138, May 2025.

[4] C. M. Perou, T. Sørlie, M. B. Eisen, and et al., “Molecular portraits of human breast tumours,” Nature, vol. 406, no. 6797, pp. 747–752, Aug. 2000.

[5] Cancer Genome Atlas Network, “Comprehensive molecular portraits of human breast tumours,” Nature, vol. 490, no. 7418, pp. 61–70, Oct. 2012.

[6] R. Callahan and S. Hurvitz, “Her2-positive breast cancer: Current management of early, advanced, and recurrent disease,” Current Opinion in Obstetrics & Gynecology, vol. 23, no. 1, pp. 37–43, Feb. 2011.

[7] D. J. Slamon, B. Leyland-Jones, S. Shak, and et al., “Use of chemotherapy plus a monoclonal antibody against her2 for metastatic breast cancer that overexpresses her2,” New England Journal of Medicine, vol. 344, no. 11, pp. 783–792, Mar. 2001.

[8] M. J. Piccart-Gebhart, M. Procter, B. Leyland-Jones et al., “Trastuzumab after adjuvant chemotherapy in HER2-positive breast cancer,” New England Journal of Medicine, vol. 353, no. 16, pp. 1659–1672, Oct. 2005.

[9] J. Baselga, J. Cortés, S.-B. Kim, and et al., “Pertuzumab plus trastuzumab plus docetaxel for metastatic breast cancer,” New England Journal of Medicine, vol. 366, no. 2, pp. 109–119, Jan. 2012.

[10] S. M. Swain, J. Baselga, S.-B. Kim, and et al., “Pertuzumab, trastuzumab, and docetaxel in her2-positive metastatic breast cancer,” New England Journal of Medicine, vol. 372, no. 8, pp. 724–734, Feb. 2015.

[11] A. N. Hata, M. J. Niederst, H. L. Archibald, M. Gomez-Caraballo, F. M. Siddiqui, H. E. Mulvey, and et al., “Overcoming resistance to tumor-targeted and immune-targeted therapies,” Clinical Cancer Research, vol. 27, no. 14, pp. 3909–3916, Jul. 2021.

[12] R. A. Burrell, N. McGranahan, J. Bartek, and C. Swanton, “The causes and consequences of genetic heterogeneity in cancer evolution,” Nature, vol. 501, pp. 338–345, 2013.

[13] A. Marusyk, M. Janiszewska, and K. Polyak, “Intratumor heterogeneity: The rosetta stone of therapy resistance,” Cancer Cell, vol. 37, pp. 471–484, 4 2020.

[14] N. McGranahan and C. Swanton, “Clonal heterogeneity and tumor evolution: Past, present, and the future,” Cell, vol. 168, no. 4, pp. 613–628, Feb. 2017.

[15] S. R. y Cajal, M. Sesé, C. Capdevila, T. Aasen, L. D. Mattos-Arruda, S. J. Diaz-Cano, J. Hernández-Losa, and J. Castellví, “Clinical implications of intratumor heterogeneity: challenges and opportunities,” Journal of Molecular Medicine, vol. 98, pp. 161–177, 2 2020.

[16] S. Shenoy, “Cell plasticity in cancer: A complex interplay of genetic, epigenetic mechanisms and tumor micro-environment,” Surgical Oncology, vol. 34, pp. 154–162, 9 2020.

[17] P. B. Gupta, C. M. Fillmore, G. Jiang, and et al., “Stochastic state transitions give rise to phenotypic equilibrium in populations of cancer cells,” Cell, vol. 146, no. 4, pp. 633–644, Aug. 2011.

[18] D. Shah and C. Osipo, “Cancer stem cells and her2 positive breast cancer: The story so far,” Genes & Diseases, vol. 3, pp. 114–123, 6 2016.

[19] N. V. Jordan, A. Bardia, B. S. Wittner, C. Benes, M. Ligorio, Y. Zheng, M. Yu, T. K. Sundaresan, J. A. Licausi, R. Desai, R. M. O’Keefe, R. Y. Ebright, M. Boukhali, S. Sil, M. L. Onozato, A. J. Iafrate, R. Kapur, D. Sgroi, D. T. Ting, M. Toner, S. Ramaswamy, W. Haas, S. Maheswaran, and D. A. Haber, “Her2 expression identifies dynamic functional states within circulating breast cancer cells,” Nature, vol. 537, pp. 102–106, 8 2016.

[20] R. Z. Granit, H. Masury, R. Condiotti, Y. Fixler, Y. Gabai, T. Glikman, S. Dalin, E. Winter, Y. Nevo, E. Carmon, T. Sella, A. Sonnenblick, T. Peretz, U. Lehmann, K. Paz, F. Piccioni, A. Regev, D. E. Root, and I. Ben-Porath, “Regulation of cellular heterogeneity and rates of symmetric and asymmetric divisions in triple-negative breast cancer,” Cell Reports, vol. 24, pp. 3237–3250, 9 2018.

[21] A. R. Freischel, M. Damaghi, J. J. Cunningham, A. Ibrahim-Hashim, R. J. Gillies, R. A. Gatenby, and J. S. Brown, “Frequency-dependent interactions determine outcome of competition between two breast cancer cell lines,” Scientific Reports 2021 11:1, vol. 11, pp. 1–18, 3 2021.

[22] A. T. Baker, A. Zlobin, and C. Osipo, “Notch–egfr/her2 bidirectional crosstalk in breast cancer,” Frontiers in Oncology, vol. 4, no. 360, 2014.

[23] K. Yao, X. Y. Zhan, M. Feng, K. F. Yang, M. S. Zhou, and H. Jia, “Furin, adam, and γ-secretase: Core regulatory targets in the notch pathway and the therapeutic potential for breast cancer,” Neoplasia, vol. 57, p. 101041, 11 2024.

[24] D. L. Abravanel, G. K. Belka, T. C. Pan, D. K. Pant, M. A. Collins, C. J. Sterner, and L. A. Chodosh, “Notch promotes recurrence of dormant tumor cells following her2/neu-targeted therapy,” Journal of Clinical Investigation, vol. 125, pp. 2484–2496, 6 2015.

[25] M. Capulli, M. Haffner-Luntzer, and et al., “Notch2 pathway mediates breast cancer cellular dormancy and mobilisation in the bone microenvironment,” British Journal of Cancer, vol. 121, no. 2, pp. 157–166, Jul. 2019.

[26] E. W. J. Mollen, J. Ient, V. C. G. Tjan-Heijnen, H. H. Boersma, L. Miele, M. L. Smidt, and M. Vooijs, “Moving breast cancer therapy up a notch,” Frontiers in Oncology, vol. 8, p. 518, Oct. 2018.

[27] X. Li and D. Thirumalai, “A mathematical model for phenotypic heterogeneity in breast cancer with implications for therapeutic strategies,” Journal of the Royal Society Interface, vol. 19, 2022.

[28] A. M. Jarrett, A. Shah, M. J. Bloom, M. T. McKenna, D. A. Hormuth, T. E. Yankeelov, and A. G. Sorace, “Experimentally-driven mathematical modeling to improve combination targeted and cytotoxic therapy for her2+ breast cancer,” Scientific Reports 2019 9:1, vol. 9, pp. 1–12, 9 2019.

[29] P. Jain, R. Kizhuttil, M. B. Nair, S. Bhatia, E. W. Thompson, J. T. George, and M. K. Jolly, “Cell-state transitions and density-dependent interactions together explain the dynamics of spontaneous epithelial-mesenchymal heterogeneity,” iScience, vol. 27, no. 7, pp. 110 310–, 2024.

[30] R. W. Holloway and P. A. Marignani, “Targeting mtor and glycolysis in her2-positive breast cancer,” Cancers, vol. 13, 6 2021.

[31] E. A. Wellberg, S. Johnson, J. Finlay-Schultz, A. S. Lewis, K. L. Terrell, C. A. Sartorius, E. D. Abel, W. J. Muller, and S. M. Anderson, “The glucose transporter glut1 is required for erbb2-induced mammary tumorigenesis,” Breast Cancer Research, vol. 18, no. 1, p. 131, 2016.

[32] L. Pan, J. Li, Q. Xu, Z. Gao, M. Yang, X. Wu, and X. Li, “Her2/pi3k/akt pathway in her2-positive breast cancer: A review,” Medicine (Baltimore), vol. 103, p. e38508, 2024.

[33] A. Rohatgi, “Webplotdigitizer,” https://automeris.io/WebPlotDigitizer, software for extracting data from plots.

[34] K. Vipparthi, K. Hari, P. Chakraborty, S. Ghosh, A. K. Patel, A. Ghosh, N. K. Biswas, R. Sharan, P. Arun, M. K. Jolly, and S. Singh, “Emergence of hybrid states of stem-like cancer cells correlates with poor prognosis in oral cancer,” iScience, vol. 25, no. 5, p. 104317, 2022.

[35] E. B. Gunnarsson, B. V. Magnússon, and J. Foo, “Optimal dosing of anti-cancer treatment under drug-induced plasticity,” NPJ systems biology and applications, vol. 11, no. 1, pp. 98–15, 2025.

[36] J. L. Gevertz, J. M. Greene, S. Prosperi, N. Comandante-Lou, and E. D. Sontag, “Understanding therapeutic tolerance through a mathematical model of drug-induced resistance,” NPJ systems biology and applications, vol. 11, no. 1, pp. 30–15, 2025.

